# Dynamics of the energy seascape can explain intra-specific variations in sea-crossing behaviour of soaring birds

**DOI:** 10.1101/733063

**Authors:** E. Nourani, W. M. G. Vansteelant, P. Byholm, K. Safi

**Affiliations:** Department of Migration, Max Planck Institute of Animal Behavior, Radolfzell, Germany; Department of Biology, University of Konstanz, Germany; Theoretical and computational Ecology, Institute for Biodiversity and Ecosystem Dynamics, University of Amsterdam, The Netherlands; Novia University of Applied Sciences, Ekenäs, Finland

**Keywords:** Energy landscape, water-crossing, thermal uplift, soaring, ecological barrier, temperature gradient

## Abstract

Thermal soaring birds extract energy from the atmosphere to achieve energetically low-cost movement. When encountering regions that are energetically costly to fly over, such as open seas, they attempt to adjust the spatio-temporal pattern of their passage to maximize energy extraction from the atmosphere over these ecological barriers. We apply the concept of energy landscapes to investigate the spatio-temporal dynamics of energy availability over the open sea for soaring flight. We specifically investigated how the “energy seascape” may shape age-specific sea-crossing behaviour of European honey buzzards *Pernis apivorus* over the Mediterranean Sea in autumn. We found uplift potential over the sea to be the main determinant of sea-crossing length, rather than wind conditions. Considering this variable as a proxy for available energy over the sea, we constructed the energy seascape for the autumn migration season using forty years of temperature data. Our results indicate that early-migrating adult buzzards are likely to encounter adverse energy subsidence over the Mediterranean, whereas late-migrating juveniles face less adverse flight conditions, and even conditions conducive to soaring flight. Our study provides evidence that the dynamics of the energy landscape can explain intra-specific variation in migratory behaviour also at sea.

## Introduction

Flying animals have evolved to interact with their highly dynamic atmospheric environments and are selected for energetic efficiency. Many species extract energy from the atmosphere by flying in suitable winds [1-3] and reducing energy expenditure by avoiding headwind and adverse weather conditions [4]. Thermal soaring birds (hereafter soaring birds) are masters of extracting energy for flight by relying on uplifts [5], which are vertically rising air currents that subsidize or entirely account for energy expenditure when soaring [6, 7]. Therefore, energy availability landscapes (see [8, 9]) for these species can be defined as a function of uplift potential and wind conditions.

Open seas and oceans are generally considered energetically costly to cross for soaring birds [10-12]. The low, or negative, uplift potential over these ecological barriers, especially in temperate regions, requires the birds to sustain long bouts of flapping flight. While their large wing area to body mass ratio makes soaring birds adapted to extract energy from uplifts, flapping flight, precisely because of the large wings, represents an expensive task [13, 14]. As such, various obligate soaring birds rely on traditional overland detours [15, 16], the cost of which is offset by their low energy consumption in thermal-soaring flight [17]. Most of the landbirds capable of thermal soaring are facultative soaring migrants. These species have a lower wing area to body mass ratio with which they can sustain long bouts of flapping flight if needed, for example over open water, as in the Eleonora’s flacon *Falco eleonorae* and the osprey *Pandion haliaetus*, [11, 18].

The morphological differences characterising obligate and facultative soaring species alone, however, cannot explain the propensity for sea-crossing, particularly when there is intra-specific variation in this behaviour. Long sea-crossings (i.e. 10s – 100s km) by soaring birds are usually associated with suitable atmospheric support. In many cases this comes in the form of supportive wind [12, 19-21]. More recently, it has also been shown to occur in the form of updrafts [18, 22]. We therefore argue that intra-specific variability in sea-crossing propensity can be explained by spatial and temporal dynamics of atmospheric energy availability over the open sea.

Here, we apply the concept of energy landscapes [8, 9] to quantify the spatio-temporal dynamics of potential energy availability over the sea for thermal soaring flight, hence using the phrase “energy seascape”. We expect that energy seascape dynamics help explain the differential sea-crossing behaviour of juvenile vs. adult European honey buzzards *Pernis apivorus* during their autumn migration from Europe to Africa. Both the propensity to cross the sea [12] and the timing of migration [23] differs between age groups in this species. We hypothesize that juveniles are able to migrate longer distances over the Mediterranean Sea due to the prevalence of more supportive atmospheric conditions during their late migration period. Wind conditions likely play a critical role in facilitating these sea-crossings. Yet, recent findings by Duriez *et al.* [18] show that soaring birds are capable of prolonged thermal soaring over the Mediterranean Sea, suggesting that uplift potential could play an underappreciated role in the sea-crossing propensity of this facultative soaring bird as well.

## Methods

We used GPS-tracking data collected for autumn migration of honey buzzards breeding in Finland. Data included complete autumn journeys for 22 juveniles [24] on their first autumn migration between 2011-2013 and nine adult birds of unknown age between 2011-2016 (ESM 1). We had data for 2-5 years for the adults, totalling 29 complete tracks (Fig. ESM 2). All data were resampled to an hourly interval to achieve consistent frequency for all tracks.

We extracted sections of the tracks that overlapped with the Mediterranean Sea and calculated its length (including islands). Each track was annotated with Julian date of first encountering the Mediterranean Sea (hereafter date of arrival at sea), wind support, crosswind, and temperature gradient over the sea (*ΔT*) as a proxy for uplift potential over water. Positive values of *ΔT* (i.e. warmer sea than air) indicate upward moving air and correspond to higher uplift potential (see [18]), while negative values indicate subsidence. Environmental data were obtained through the Env-Data track annotation service of Movebank [25].

To understand whether atmospheric conditions affect the risk birds were willing to take when crossing the sea, we modelled the length of the journeys the birds took when crossing the Mediterranean as a function of wind support, cross-wind, and *ΔT* along the track, as well as date of arrival at sea, using a linear model with a random intercept effect for year. We did not include age group as an explanatory variable in the model, because of its high correlation with date of arrival at sea (i.e. adults arrived at sea earlier than juveniles).

We then constructed energy seascapes using the most influential variable in our linear model. To further investigate the spatio-temporal dynamics of energy seascapes, we predicted the most influential variable from the linear model as a function of latitude, longitude, and time, in a generalized additive model (GAM; [26, 27]), using 40 years of data (1979-2018) for the autumn season (August-October). To produce energy seascape maps, we used the model to make predictions for two time periods corresponding to juvenile (Sep. 25-Oct. 23) and adult (Aug 30-Oct. 4) migration over the sea. This data was a subset of the 40-year dataset that was used to build the model. To create smooth maps, we spatially interpolated the output raster layers to 1-km resolution (See ESM 3 for more detail on methodology).

## Results

We identified temperature gradient, delta T (a proxy for uplift potential), as the most important atmospheric variable contributing to sea-crossing distance. Our best model (defined by the highest deviance explained; model C in Table 1) showed longer sea-crossing distance in migrating honey buzzards occur with an increasing temperature gradient (warmer sea surface temperature than air). We also found a considerable effect of crosswind and date of arrival at sea (Table 1).

**Table 1.** Results of linear modelling with a random effect for year. Distance of sea-crossing over the Mediterranean Sea was investigated as a function of wind support, crosswind, temperature gradient, and date of arrival at sea.

| Model | Coefficients (SE) |  |  |  |  | Random effects | AIC | Adjusted R<sup>2</sup> |
| --- | --- | --- | --- | --- | --- | --- | --- | --- |
| | Intercept | Wind support | Crosswind | $\Delta T$ | Julian date | | | |
| A | 805.69 (33.69) | 2.21 (26.88) | 58.82 (26.01) | - | 51.04 (26.81) | Year<sup>1</sup> | 576.21 | 0.20 |
| B | 806.36 (31.96) | - | - | 71.61 (26.39) | 30.17 (27.32) | Year<sup>2</sup> | 580.98 | 0.23 |
| C | 803.58 (28.66) | - | 56.77 (23.30) | 69.22 (25.01) | 20.44 (25.96) | Year<sup>3</sup> | 569.20 | 0.33 |

<sup>1</sup>Between-year standard deviation: 49.82; residual standard deviation: 162.90
<sup>2</sup>Between-year standard deviation: 46.15; residual standard deviation: 158.50
<sup>3</sup>Between-year standard deviation: 37.75; residual standard deviation: 150.93

Temperature gradient was then used in a GAM to construct energy seascapes. The importance of timing and temperature gradient over the sea during the migration of juveniles was confirmed by the results of this analysis (Table 2). These results showed that as the autumn migration season progresses, delta T increases (Figs. 1 & 2). Energy seascape maps over the Mediterranean Sea showed that, earlier in autumn, when adults arrive at the sea, delta T is low or negative, causing subsidence (Fig. 1a). Later in the season, however, a warmer sea surface compared to the air above it provides uplift potential (Fig. 1b).

**Table 2.**
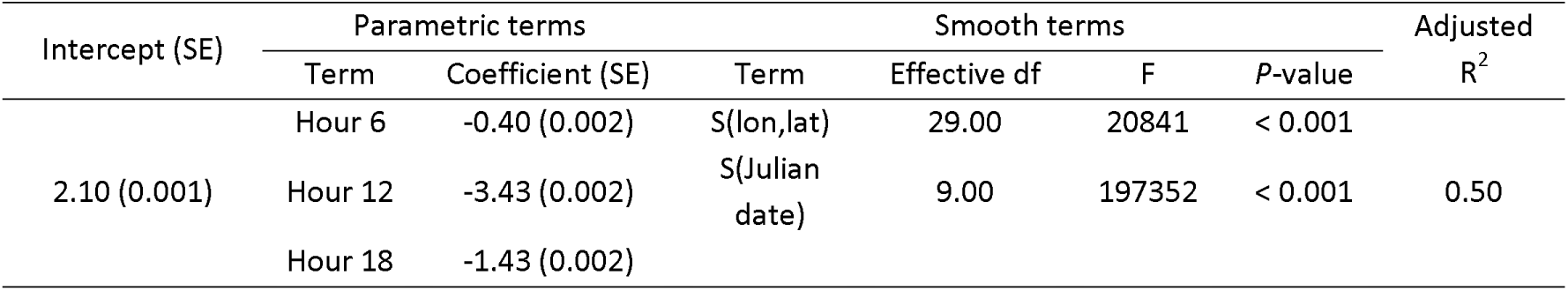
Results of generalized additive model predicting temperature gradient as a function of longitude, latitude, Julian date, and hour. The model includes a 2-dimensional smooth term for longitude and latitude, a smooth term for Julian date, and a parametric term for hour as a factor.

**Figure 1.**
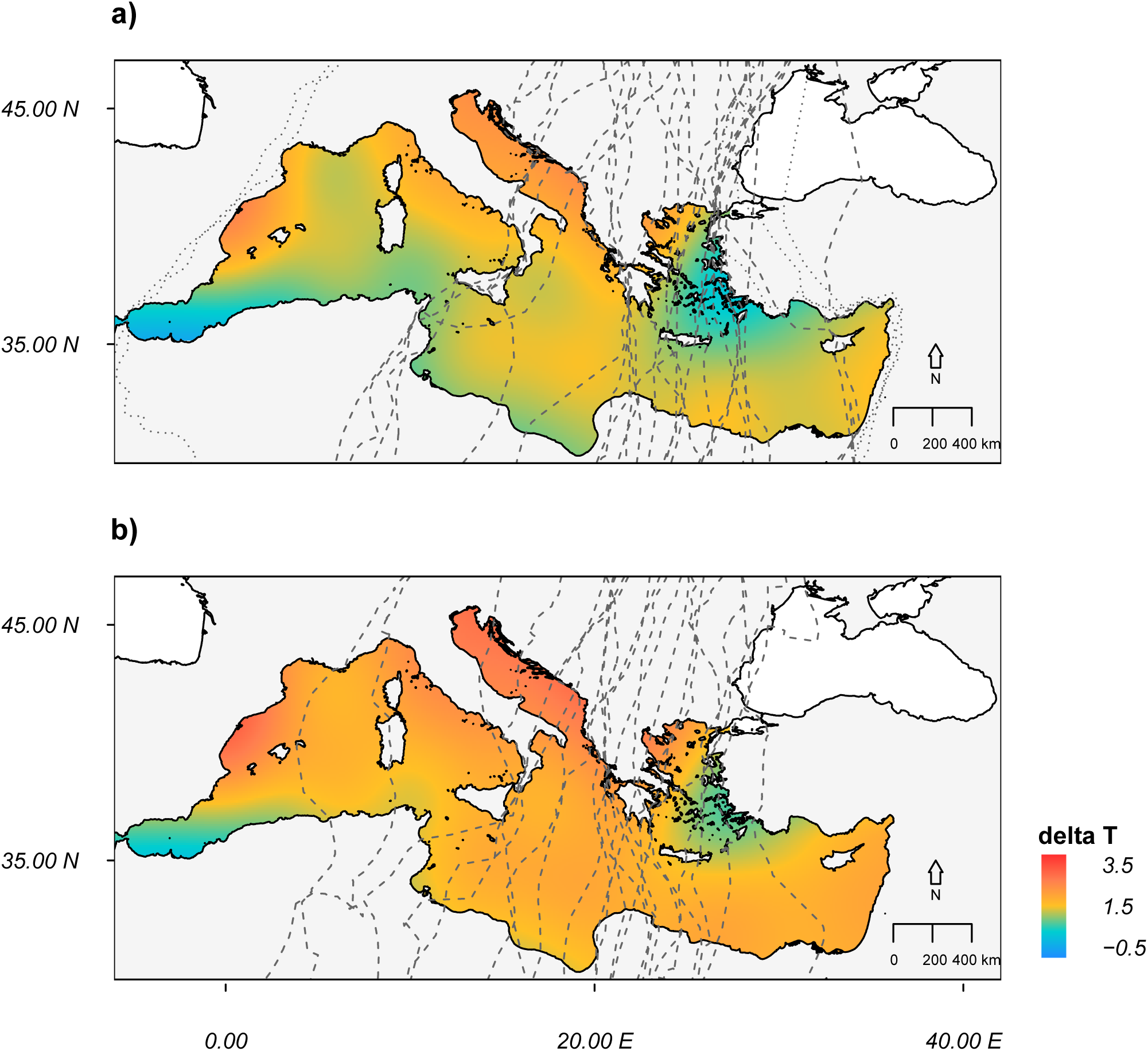
Energy seascapes constructed by predicting temperature gradient (delta T; °C) over the Mediterranean Sea for the period of honey buzzard autumn migration for a) adults (Aug 30-Oct. 4) and b) juveniles (Sep. 25-Oct. 23) over the sea. The dashed lines in each figure show autumn migration tracks for the corresponding age group. The dotted lines in (a) correspond to migration tracks removed from the linear model analysis (For complete migration tracks within the flyway, see Fig. ESM 2).

**Figure 2.**
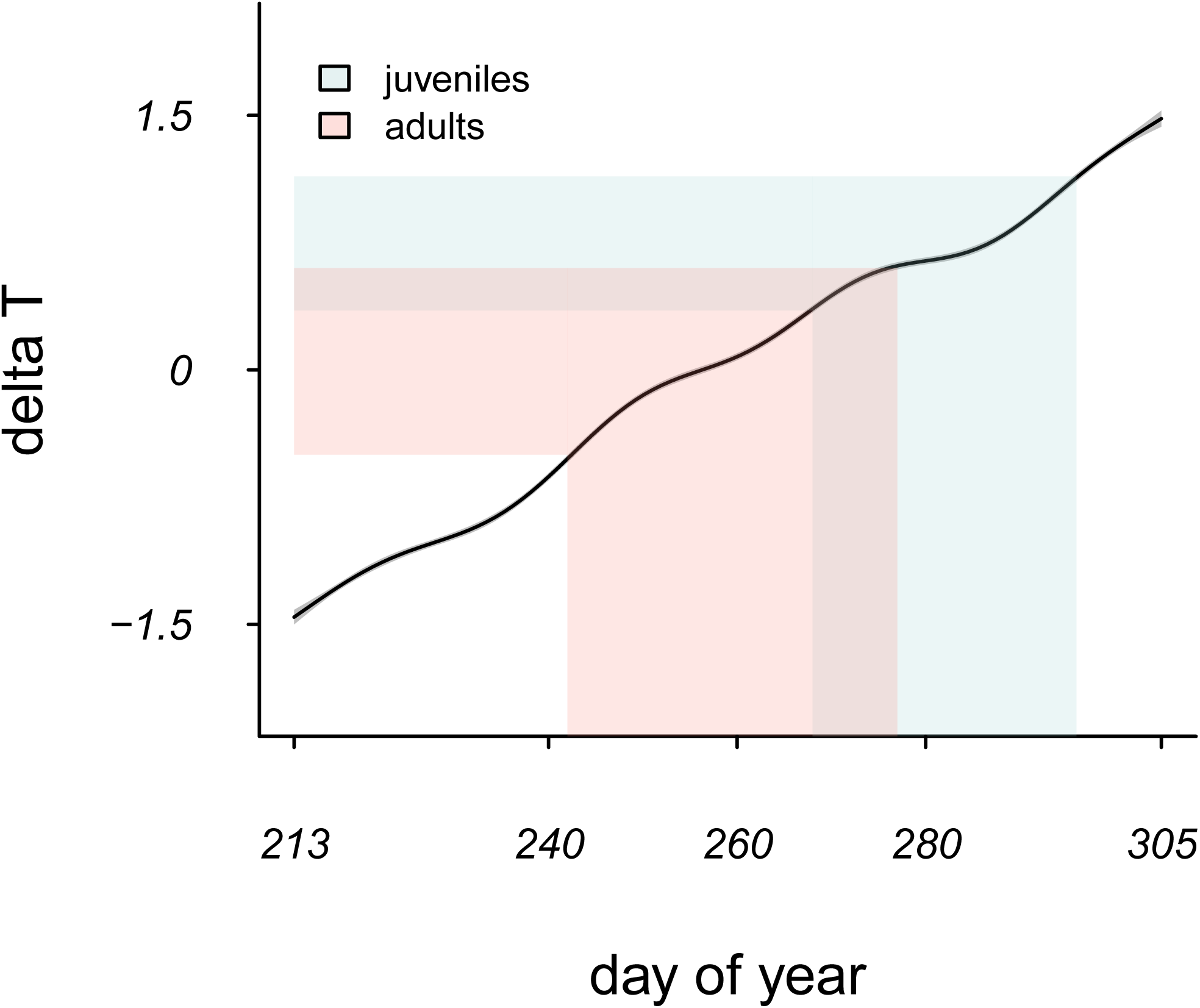
Smooth term showing the relationship between temperature gradient (delta T) and Julian Date over the Mediterranean Sea. The pink and blue shaded areas corresponds to the periods of adult and juvenile migration over the sea, respectively.

## Discussion

We constructed energy seascapes using temperature gradient, a proxy for uplift potential, that was recently shown by Duriez *et al.* [18] to be correlated with soaring flight over the sea. Our approach to quantifying the energy seascape using forty years of temperature data demonstrated a consistent pattern of increasing uplift potential as autumn migration season progresses. This finding can be directly applied to other studies of bird migration over the Mediterranean Sea in autumn. In a recent study of facultative sea-crossing in white storks *Ciconia ciconia* for example, Becciu *et al.* [28] found a positive relationship between Julian date and the propensity of sea-crossing in autumn. This pattern could be related to the temporally dynamic uplift potential over the Mediterranean Sea that simply makes sea-crossing energetically cheaper later in autumn. For studies focusing on sea-crossing in other temporal and/or spatial contexts as we describe here, our methodology can be applied for constructing energy seascapes to explore the dynamics of energy availability over the open sea.

Bird flight over open water is often facilitated by wind support [19-21, 29], a variable that has great potential to be used for constructing energy seascapes. Nourani *et al.* [20] found that seasonality in wind conditions is responsible for the seasonal sea-crossing strategies in Oriental honey buzzards *P. ptilorhynchus* in East Asia. Wind support has a dominant influence on the hourly and daily migration performance of adult European honey buzzards [30], and large-scale wind regimes seem to mould seasonal loop migrations of adult honey buzzards at the flyway-scale [31]. In our current study, both age groups crossed the sea with comparably moderate wind support (ESM 4).

We found that uplift potential over the Mediterranean Sea can account for the variation in the length of sea-crossing performed by honey buzzards. By quantifying the temporal dynamics in uplift potential in energy seascape maps, we further showed that flight over the sea is energetically cheaper later in the autumn season when juvenile birds are migrating. We do not suggest that this is the causal mechanism of differential migration onset between the two age groups. Indeed, earlier migration timing in adults is likely to be related to carry-over effects that a delay in autumn migration could have on the later stages of their life-cycle, such as failure to secure prey-rich wintering territories [32]. Later migration onset in juveniles might be related to the time required for their development to become capable of migration [33]. Nevertheless, our results indicate that crossing the Mediterranean Sea late in the autumn is less energetically demanding than commonly assumed in the literature. This may be a driver of how such age-specific variation in the timing of migration and the propensity to cross the sea has been selected for and maintained in the honey buzzard population.

While first-year migrant birds generally suffer high mortality [34, 35] and juvenile soaring migrants are unlikely to survive long sea-crossings [36, 37], juvenile honey buzzards have high survival while migrating over the Mediterranean Sea [24]. Juvenile honey buzzards on their first autumn migration travel alone or in small flocks and are likely to have inferior soaring abilities compared to adults. Following a general southward direction of movement, they are assisted by the wind [24]. Whenever they encounter the Mediterranean Sea, they cross without suffering high mortality rates [24]. Yet the birds’ route selection criteria eventually do become more sophisticated with age and experience [38], likely improving the use of the wind and uplift conditions [39, 40]. Along with the higher cost of flight over the sea during early autumn, these improvements can explain why adults eventually prefer to reduce sea-crossing. Integrating wind support as well as uplift potential in energy landscapes at a flyway-scale may help us quantify the costs and benefits of alternative schedules and routes for migration, and thus to explain age-specific as well as population-specific sea-crossing behaviours and other aspects of migration strategies in facultative soaring migrants.

## Conclusion

By quantifying energy seascapes using temperature gradient over the sea as a proxy for thermal soaring opportunities, we confirm that, contrary to the traditional beliefs about the impossibility of soaring flight over temperate seas, uplift potential over the Mediterranean Sea can aid terrestrial birds in tackling this ecological barrier. Additionally, we suggest that the dynamics of the energy seascape might be a driver for the maintenance of variation in sea-crossing behaviour. Thus, our study provides first evidence that the dynamics of the energy landscape can explain intra-specific variations in migratory behaviour.

## Supporting information

ESM 1

ESM 2

ESM 3

ESM 4

## Acknowledgements

We would like to thank G. Fehlmann, M. Panuccio, B. Klump, and M. Chimento for their comments on the manuscript. We thank M. Honkiniemi, A. Rossi, A. Rantamäki, J. Valkama, J. Kivelä, I. Nousiainen, K. Palo and M. Lehtonen for assisting with fieldwork in Finland. We acknowledge the role of B. U. Meyburg in the initiation of this collaboration.

