## Supplementary figures and images for "Dynamics of the energy seascape can explain intra-specific variations in sea-crossing behaviour of soaring birds"

### ESM 2

#### Autumn migration of European honey buzzards from Finland

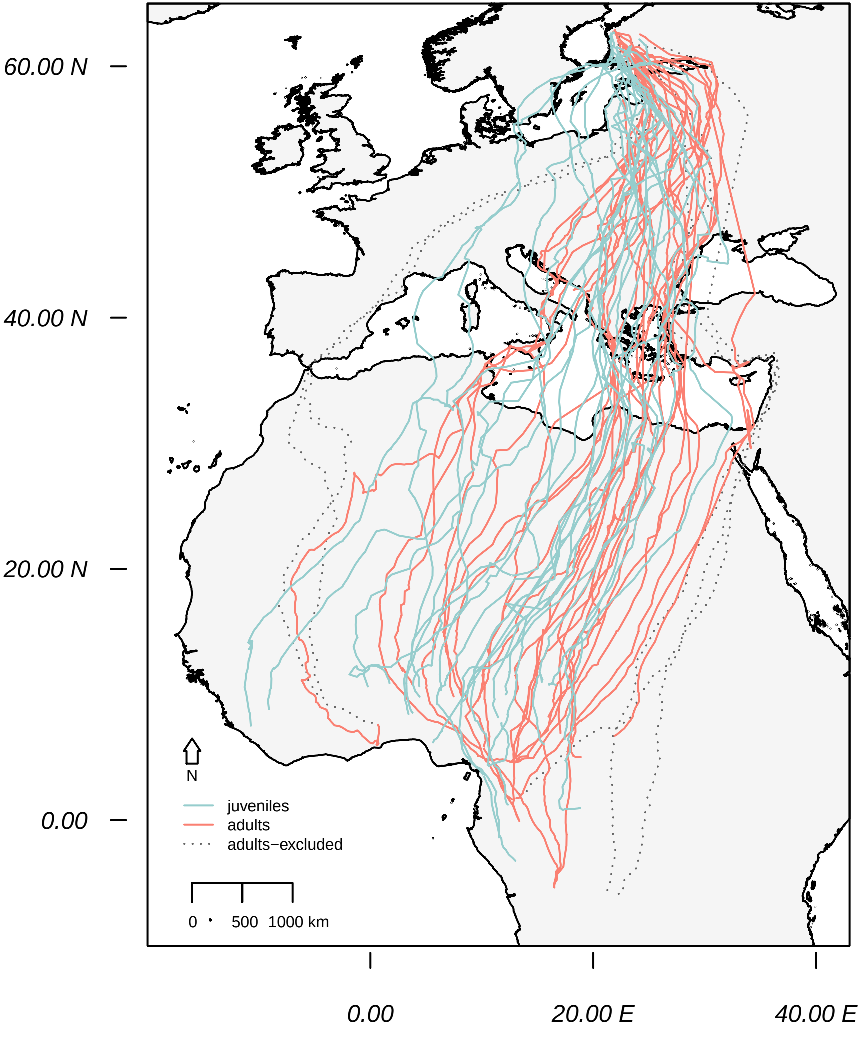

### ESM 4

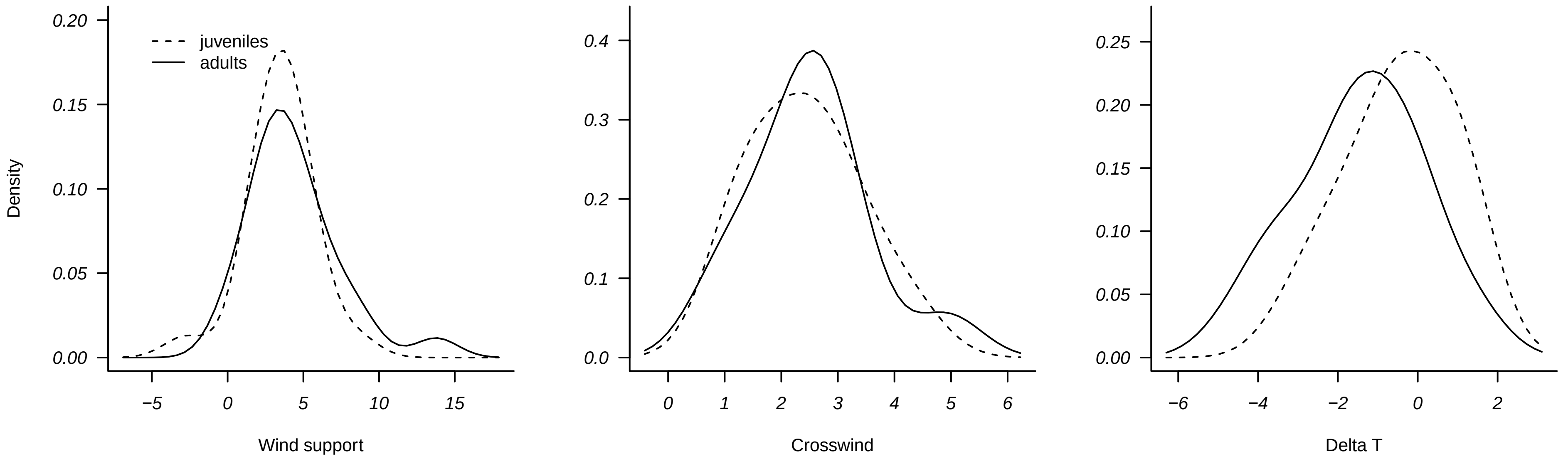
