## Supplementary material for "Dynamics of the energy seascape can explain intra-specific variations in sea-crossing behaviour of soaring birds": ESM 3

### ESM 3. Details of Methodology

#### *Tracking data*

We used GPS-tracking data collected for autumn migration of honey buzzards breeding in Finland. Data included complete autumn journeys for 22 juveniles [1] on their first autumn migration over 2011-2013 and nine adult birds of unknown age over 2011-2016 (ESM 1). We had data over 2-5 years for the adults, totalling 29 complete tracks (Fig. ESM 2). The amount and type of data varied among tags depending on the devices and programming schedule (see [1] and ESM 1), but all data was resampled to hourly or larger intervals due to gaps in data or lower measurement resolution ( e.g. at night and in the tropical forest).

For the purpose of this study, we extracted sections of the tracks that overlapped with the Mediterranean Sea. To prevent over-dispersion in our analyses, we removed tracks with shorter water-crossing lengths than the 10% quantile of the lengths of all water-crossing tracks (415.18 km). These tracks (Figs. 1 & ESM 2) corresponded to journeys of adult birds that clearly avoided the Mediterranean Sea by flying through the strait of Gibraltar (n = 2) or the eastern flyway (n = 3).

#### *Track annotation*

Each sea-crossing track was annotated with temporal and environmental variables. We calculated the length of the sea-crossing track (including islands) and the Julian date of first encountering the Mediterranean Sea (hereafter date of arrival at sea). Every sea-crossing track (excluding islands) was interpolated with regular points 500 m apart and annotated using the Env-Data track annotation service of Movebank [2]. For each point, eastward (u) and northward (v) components of the wind (m/s) at 925 hPa pressure level, sea surface temperature (K), and temperature at 2-m above ground (K) were obtained with 0.7° resolution (all from European Centre for Medium-Range Weather Forecast; ECMWF). We selected bilinear and nearest-neighbour interpolation methods for wind and temperature, respectively. For each sea-crossing track, we calculated wind support (wind parallel to flight direction) and crosswind (wind perpendicular to flight direction) from one point to the next and calculated averages for each track. Average crosswind was calculated as the mean of absolute values along each track, as we were not interested in the direction of crosswind, but its overall strength. For each point, we calculated temperature gradient ($\Delta T$) as the difference between sea surface temperature and temperature at 2 m above ground and averaged all $\Delta T$ values along each track. We did not convert the unit of temperature variables from K to °C, because the two units have the same magnitude. Hence, *ΔT* values that we calculated can be interpreted in either unit. We used *ΔT* as an indicator of uplift potential over water. Positive values of *ΔT* (i.e. warmer sea than air) indicate upward moving air and correspond to higher uplift potential (see [3]), while negative values indicate subsidence.

*Statistical analysis*

To understand whether atmospheric conditions facilitate sea-crossing, we investigated the distance of sea-crossing (i.e. length of the sea-crossing tracks) as a function of wind support, cross-wind, and *ΔT* along the track, as well as date of arrival at sea. The data was analysed using a linear model with a random effect for year. We did not include a random effect of individuals, because we did not have repeated tracks for most of them. Age group was also not included, because of its high correlation with date of arrival at sea (i.e. all adults arrived at sea earlier than juveniles). All fixed effects were checked for multi-collinearity and were z-transformed to ensure comparability of effect sizes. Due to the small sample size ($n = 46$), we kept the model simple by estimating a small number of parameters using the data and including no interaction terms. We built separate models for wind (Model A) and *ΔT* (Model B), and a model combining the variables with highest effect sizes in the previous two models (Model C). Models were compared based on the amount of variation that they explained.

We then constructed energy seascapes using the most influential variable in our linear model (delta T). To further investigate the temporal and spatial dynamics of energy seascapes, we predicted this variable as a function of latitude, longitude, and time, in a generalized additive model (GAM; [4]). To do this, we obtained the relevant atmospheric data for the Mediterranean Sea for autumn (August- October) from ECMWF ERA-Interim reanalysis archive (*https://www.ecmwf.int/en/forecasts/datasets/reanalysis-datasets/era-interim*) for all the years for which data was available (1979-2018). We included a two-dimensional smoothing term for latitude and longitude and a one-dimensional smoothing term for Julian date using penalized regression splines. ERA-Interim data are provided for every six hours. We therefore included time of the day (00:00, 06:00, 12:00, and 18:00 UTC) as a factor variable to control for this variation. We used the mgcv package [5] for building the GAM and used a generalized cross validation criterion to estimate the best smoothing parameter. To produce energy seascape maps, we used the model to make predictions for two time periods corresponding to juvenile (Sep. 25-Oct. 23) and adult (Aug 30-Oct. 4) migration over the sea. This data was a subset of the 40-year dataset that was used to build the model. To create smooth maps, we spatially interpolated the output raster layers to 1-km resolution.

*References*

[1] Vansteelant, W. M. G., Kekkonen, J., Byholm, P. 2017 Wind conditions and geography shape the first outbound migration of juvenile honey buzzards and their distribution across sub-Saharan Africa. *Proceedings of the Royal Society B: Biological Sciences*. **284**, (doi: 10.1098/rspb.2017.0387)

[2] Dodge, S., Bohrer, G., Weinzierl, R., Davidson, S. C., Kays, R., Douglas, D. 2013 The environmental-data automated track annotation (Env-DATA) system: linking animal tracks with environmental data. *Movement Ecology*. **1**, (doi: 10.1186/2051-3933-1-3)

[3] Duriez, O., Peron, G., Gremillet, D., Sforzi, A., Monti, F. 2018 Migrating ospreys use thermal uplift over the open sea. *Biology Letters*. **14**, 20180687. (doi: 10.1098/rsbl.2018.0687)

[4] Wood, S. N. 2017 *Generalized additive models: an introduction with R*. Chapman and Hall/CRC.

[5] Wood, S. Package 'mgcv'. R package version v.1.8-28. 2019.
